# The developmental genetic architecture of vocabulary skills during the first three years of life: Capturing emerging associations with later-life reading and cognition

**DOI:** 10.1101/2020.10.05.325993

**Authors:** Ellen Verhoef, Chin Yang Shapland, E. Fisher Simon, Philip S. Dale, Beate St Pourcain

## Abstract

Individual differences in early-life vocabulary measures are heritable and associated with subsequent reading and cognitive abilities, although the underlying mechanisms are little understood. Here, we (i) investigate the developmental genetic architecture of expressive and receptive vocabulary in toddlerhood and (ii) assess origin and developmental stage of emerging genetic associations with mid-childhood verbal and non-verbal skills.

Studying up to 6,524 unrelated children from the population-based Avon Longitudinal Study of Parents and Children (ALSPAC) cohort, we dissected the phenotypic variance of longitudinally assessed early-life vocabulary measures (15-38 months) and later-life reading and cognitive skills (7-8 years) into genetic and residual components, by fitting multivariate structural equation models to genome-wide genetic-relationship matrices.

Our findings show that the genetic architecture of early-life vocabulary is dynamic, involving multiple distinct genetic factors. Two of them are developmentally stable and contribute to genetic variation in mid-childhood skills: Genetic links with later-life verbal abilities (reading, verbal intelligence) emerged with expressive vocabulary at 24 months. The underlying genetic factor explained 10.1% variation (path coefficient: 0.32(SE=0.06)) in early language, but also 6.4% (path coefficient: 0.25(SE=0.12)) and 17.9% (path coefficient: 0.42(SE=0.13)) variation in mid-childhood reading and verbal intelligence, respectively. An independent stable genetic factor was identified for receptive vocabulary at 38 months, explaining 2.1% (path coefficient: 0.15(SE=0.07)) phenotypic variation. This genetic factor was also linked to both verbal and non-verbal cognitive abilities in mid-childhood, accounting for 24.7% of the variation in non-verbal intelligence (path coefficient: 0.50(SE=0.08)), 33.0% in reading (path coefficient: 0.57(SE=0.07)) and 36.1% in verbal intelligence (path coefficient: 0.60(0.10)), corresponding to the majority of genetic variance (≥66.4%).

Thus, the genetic foundations of mid-childhood reading and cognition are diverse. They involve at least two independent genetic factors that emerge at different developmental stages during early language development and may implicate differences in cognitive processes that are already detectable during toddlerhood.

**Author summary:** Differences in the number of words young children produce (expressive vocabulary) and understand (receptive vocabulary) can be partially explained by genetic factors, and are related to reading and cognitive abilities later in life. Here, we studied genetic influences underlying word production and understanding during early development (15-38 months) and their genetic relationship with mid-childhood reading and cognitive skills (7-8 years), based on longitudinal phenotype measures and genome-wide genetic data from up to 6,524 unrelated children. We showed that vocabulary skills assessed at different stages during early development are influenced by distinct genetic factors, two of which also contribute to genetic variation in mid-childhood skills, suggesting developmental stability: Genetic sources emerging for word production skills at 24 months were linked to subsequent verbal abilities, including mid-childhood reading and verbal intelligence performance. A further independent genetic factor was identified that related to word comprehension at 38 months and also contributed to variation in later verbal as well as non-verbal abilities during mid-childhood. Thus, the genetic foundations of mid-childhood reading and cognition involve at least two independent genetic factors that emerge during early-life langauge development and may implicate differences in overarching cognitive mechanisms.

## Introduction

The number of words produced and understood by children during the first few years of life is a rapidly changing developmental phenotype that is often used to assess the level of language acquisition (1). One of the first precursors of expressive vocabulary (i.e. word production) in typically developing children is canonical babbling, which emerges around the age of four to six months (2), followed by the spontaneous production of first words between 10 to 15 months of age (3). With progressing development, the number of produced words increases, reaching a median of 40 words at 16 months (1), often trailed by a period of rapid growth till the age of about 22 months (4) and a steady increase after that. This results in the production of approximately 500 words at 30 months (5) and about 2,600 words at six years of age (6). The development of receptive vocabulary (i.e. word comprehension) typically precedes expressive vocabulary in developing children (7), with the understanding of the first few words emerging between 6 to 9 months of age (8). Thus, receptive vocabulary is often larger than expressive vocabulary in size (7). For example, the number of words understood by infants at 16 months of age has a median of 169 words, and is, thus, approximately 129 words larger compared to their expressive vocabulary at the same time (1). This discrepancy increases during development, with a receptive vocabulary size of about 20,000 to 24,000 words at the age of six years, which is about six times larger than its expressive counterpart (6).

The rate of language acquisition, and thus vocabulary size, varies between children during early language development (9,10). These large interindividual differences can partially be explained by genetic variation. Twin studies estimated that genetic influences could account for 17% to 25% of variation in expressive vocabulary at 24 months (11,12), 10% to 14% of variation in expressive vocabulary at 36 months (11) and 28% of variation in receptive vocabulary at 14 months (13). Studies using genotype data from unrelated children provided similar estimates, with single-nucleotide polymorphism heritability (SNP-h^2^) estimates of 13% to 14% for expressive vocabulary at 15 to 30 months of age (14) and 12% for receptive vocabulary at 38 months of age (15).

Despite some stable genetic contributions during early development, there is evidence for age-specific genetic influences on vocabulary skills. For example, measures of expressive vocabulary size assessed between 15 and 36 months were genetically only moderately correlated, with estimates ranging from 0.48 to 0.69 (11,14). Additionally, a considerable proportion (3% to 28%) of the total variation in early expressive language assessed at 24, 36 and 48 months could be explained by measure-specific additive genetic variance and not by a shared latent factor (16). However, the field is still missing an in-depth characterisation of the genetic architecture underlying early-life vocabulary development that characterises age-specific genetic influences across infancy and toddlerhood starting from the first-word stage as well as differences between receptive and expressive language skills.

Genetic links between early language processes (assessed from 24 to 48 months of age) and subsequent language- and literacy-related abilities (assessed from mid-childhood to adolescence) have been reported by studies of both twins and unrelated individuals (15–17). This research suggested that genetic variance in mid-childhood/adolescent language, literacy and cognitive development can already be captured by genetic factors contributing to language skills in toddlerhood, i.e. before the age of four years. More specifically, genetic influences underlying receptive vocabulary at 38 months could capture, through amplification, the majority of genetic variation contributing to a wide spectrum of mid-childhood/early-adolescent literacy and (verbal) cognitive skills in a sample of unrelated individuals (15). So far, however, our understanding of the developmental origin of these factors is incomplete.

Here, we (i) examine stability and change in the developmental genetic architecture of language during the first three years of life and (ii) assess origin and developmental stage of emerging genetic associations with verbal and non-verbal abilities during mid-childhood. We model multivariate genetic architectures underlying these traits as directly captured by genome-wide information (based on genetic-relationship-matrices, GRMs) for up to 6,524 unrelated youth from the UK Avon Longitudinal Study of Parents and Children (ALSPAC) birth cohort (18,19). We apply GRM structural equation modelling (GSEM) (20), analogous to twin research-modelling techniques, and dissect the phenotypic variation into additive genetic and residual variance structures.

## Results

### Analysis strategy

A two-stage analysis strategy was followed: During the first stage of the analysis (Stage 1), we examine the multivariate genetic variance structure of expressive and receptive vocabulary from 15 to 38 months of age (Table 1). A structural equation model (SEM) only was fitted to vocabulary measures with at least nominal evidence for SNP-h^2^ (*P*<0.05). During the second stage (Stage 2), we extend these models, and assess the emerging genetic links between early-life vocabulary (15 to 38 months) and reading, verbal intelligence quotient (VIQ) scores and performance (non-verbal) intelligence scores (PIQ) during mid-childhood (7 to 8 years of age, S1 Table). For all SEMs studied, we report path coefficients (the square root of individual factor variance contributions) and the corresponding percentage of explained phenotypic variance, in addition to total SNP-h^2^, genetic and residual correlations, factorial co-heritability (the proportion of total SNP-h^2^ explained by a specific genetic factor) and bivariate heritability (the contribution of genetic factors to the observed phenotypic correlation between two measures) (S3, S4, S5 Appendix).

**Table 1.** Early-life expressive and receptive vocabulary in ALSPAC.

| Measure | Psychological instrument | Mean Score (SE) | Mean Age (SE) | N (%male) | GCTA-SNP-$h^2$ (SE) |
| --- | --- | --- | --- | --- | --- |
| Expressive vocabulary | MacArthur CDI<sup>a</sup> | 14.29 (17.76) | 1.28 (0.08) | 6,524 (51.1) | 0.11(0.05) |
|  | MacArthur CDI<sup>b</sup> | 64.21 (35.11) | 2.03 (0.09) | 6,014 (51.7) | 0.16(0.06) |
|  | MacArthur CDI<sup>b</sup> | 113.33 (17.44) | 3.21 (0.10) | 6,092 (51.4) | 0.18(0.06) |
| Receptive vocabulary | MacArthur CDI<sup>a</sup> | 75.85(31.78) | 1.28 (0.08) | 6,524 (51.1) | 0.08(0.05) |
|  | MacArthur CDI<sup>b</sup> | 109.75 (23.75) | 3.21 (0.10) | 6,092 (51.4) | 0.12(0.06) |

Expressive vocabulary and receptive vocabulary were assessed between 15-38 months of age in independent children (genetic relationship $<0.05$ ). <sup>a</sup>Adapted form of the MacArthur CDI:Words & Gestures, consisting of 134 words. <sup>b</sup>Adapted from of the MacArthur CDI:Words & Sentences, consisting of 123 words. Abbreviations: ALSPAC, Avon Longitudinal Study of Parents and Children; CDI, Communicative Development Inventory; GCTA, Genome-wide Complex Trait Analysis; $h^2$ , heritability; SNP, single-nucleotide polymorphism.

### Stage 1: The developmental genetic architecture of early-life vocabulary skills

#### Univariate SNP-heritability estimates for early-life vocabulary measures

Measures of early-life language included expressive vocabulary at 15, 24 and 38 months and receptive vocabulary at 15 and 38 months (Table 1). They were assessed with parent-reported questionnaires and analysed as rank-transformed scores (see Methods). For comparison with multivariate models, we first estimated SNP-h^2^ using Genome-based Restricted Maximum Likelihood as implemented in Genome-wide Complex Trait Analysis (GCTA) software (21). Common genetic variation accounted for a modest proportion of phenotypic variation in early-life vocabulary throughout, except for receptive vocabulary at 15 months, where SNP-h^2^ was consistent with zero (Table 1). GCTA-SNP-h^2^ estimates for expressive vocabulary at 15, 24 and 38 months were 11%(SE=5%), 16%(SE=6%) and 18%(SE=6%), respectively. For receptive vocabulary at 15 and 38 months, SNP-h^2^ was estimated at 8%(SE=5%) and 12%(SE=6%), respectively. Given little evidence for SNP-h^2^ for receptive vocabulary at 15 months (*P*>0.05; Table 1), we excluded this measure from further correlation and GSEM analyses to facilitate the convergence of the models. Note that it was not possible to include the receptive vocabulary score at 24 months due to discrepancies in the questionnaire coding scheme (see Methods).

#### Bivariate phenotypic and genetic correlations among early-life vocabulary measures

Early-life vocabulary measures were phenotypically interrelated, although correlations decreased with increasing age windows (Fig 1a). The largest phenotypic correlation (r_p_) was estimated between expressive and receptive vocabulary at 38 months (r_p_=0.63). Bivariate genetic correlations (r_g_) among early-life vocabulary measures emerged from 24 months of age onwards (Figs 1b). Mirroring phenotypic relationships, the largest genetic correlation was observed between expressive and receptive vocabulary assessed at 38 months (GCTA-r_g_=0.86(SE=0.15), *P*=0.004).

**Fig 1.**
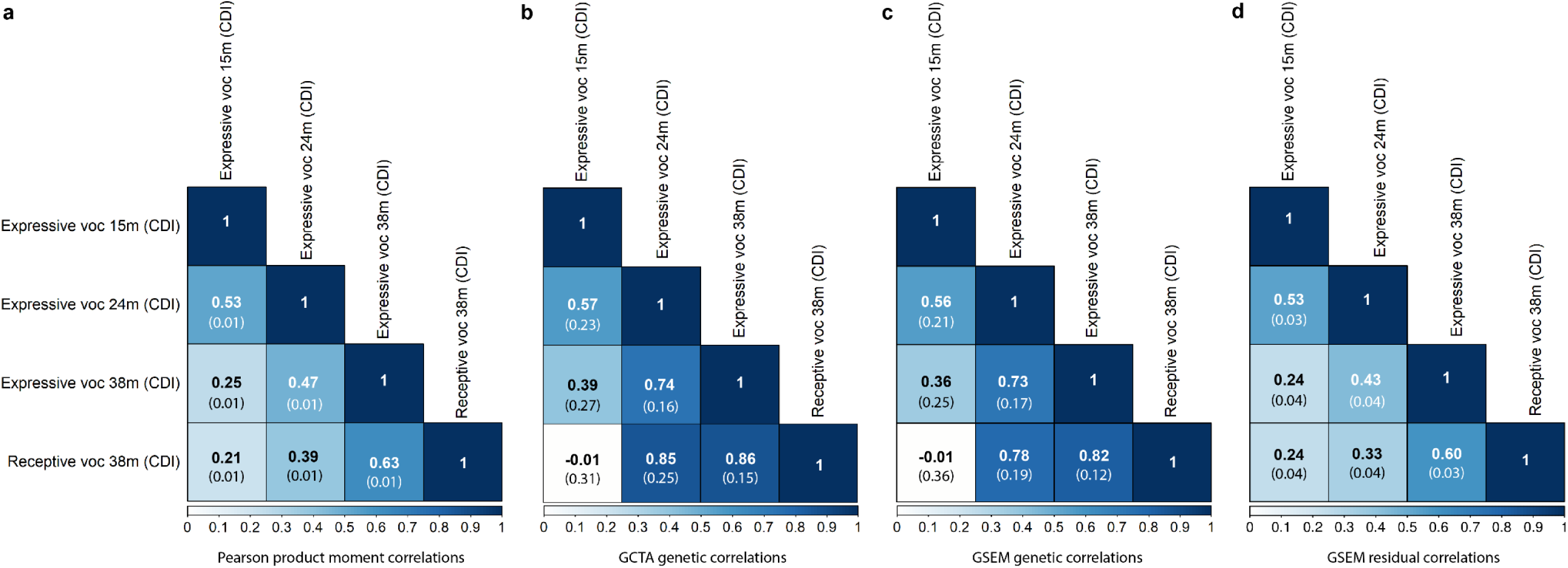
Phenotypic, genetic and residual correlations among early-life vocabulary scores (15 to 38 months). Correlation patterns are shown for rank-transformed measures with sufficient evidence for SNP-h^2^ (*P*<0.05) Standard errors are shown in brackets. **(a)** Phenotypic correlations were estimated with Pearson correlation coefficients. **(b)** GCTA genetic correlations based on GREML. **(c)** GSEM genetic correlations. **(d)** GSEM residual correlations. Abbreviations: CDI, Communicative Development Inventory; GCTA, Genome-based Restricted Maximum Likelihood as implemented in genome-wide complex trait analysis (GCTA) software; GREML, Genome-based restricted maximum likelihood; GSEM, genetic-relationship-matrix structural equation modelling; m, months; voc, vocabulary. Genetic analyses were conducted using genetic relationship matrices based on directly genotyped SNPs and individuals with a genetic relationship of <0.05

#### Multivariate genetic variance structures between early-life vocabulary measures

Using GSEM, we studied the multivariate genetic architecture underlying early vocabulary development, while allowing for both shared (i.e. across age and/or ability) and unique (i.e. age- and ability-specific) genetic influences. A multivariate SEM was fitted to expressive vocabulary at 15, 24 and 38 months as well as receptive vocabulary at 38 months (in this order), following a Cholesky decomposition. SNP-h^2^ estimates were nearly identical for all early-life vocabulary measures using univariate GCTA and multivariate GSEM approaches (Table S2). Estimated bivariate genetic correlations using GSEM were also highly consistent with GCTA findings (Figs 1b and 1c), with overlapping 95%-confidence intervals (95%-CIs). GSEM-estimated residual correlations among vocabulary measures were modest to moderate (Fig 1d), suggesting further shared aetiological mechanisms not captured by common variation.

Structural models of vocabulary measures assessed during the first three years of life revealed that the underlying genetic architecture is dynamic, with evidence for age-specific genetic influences (Fig 2). The first genetic factor (A1) accounted for 10.6%(SE=5.0%) of the phenotypic variation in expressive vocabulary at 15 months (Fig 2, S3 Table), which can be estimated by squaring the corresponding estimated path coefficient, here a_11_ (path coefficient a_11_:0.33(SE=0.08), *P*=2×10^−5^). By structural model design, the phenotypic variance explained by a_11_ corresponds to the SNP-h^2^ of expressive vocabulary at 15 months (S2 and S3 Tables). Genetic factor A1 was also related to expressive vocabulary at 24 months (path coefficient a_21_:0.21(SE=0.10), *P*=0.04), explaining 4.6%(SE=4.4%) of the phenotypic variation and accounting for almost a third of the SNP-h^2^ (factorial co-heritability: 31.2%(SE=23.4%), S4 Table). However, there was little evidence for shared genetic influences between expressive vocabulary at 15 months and either expressive or receptive vocabulary scores at 38 months (Fig 2, S3 Table). This pattern of findings suggests that genetic influences underlying expressive vocabulary at 15 months play a decreasing role during the course of later vocabulary development, consistent with data from genetic correlation and bivariate heritability analyses (Figs 1b and 1c, S5 Table).

**Fig 2.**
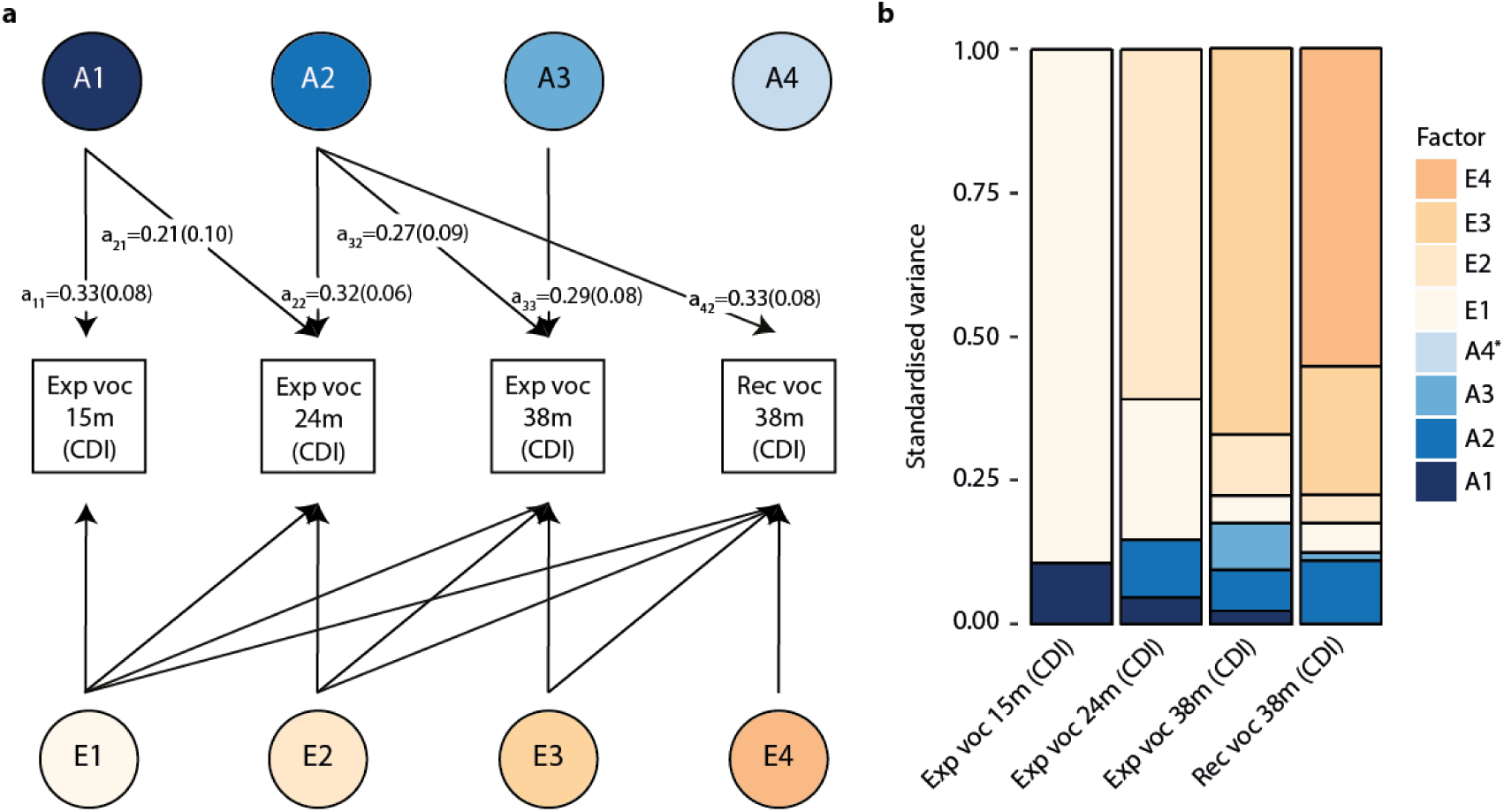
Structural model of early-life vocabulary scores (15 to 38 months) Genetic-relationship matrix structural equation modelling (GSEM) of early-life vocabulary scores (15, 24 and 38 months of age) based on all available observations for children across development (N≤6,524; Cholesky decomposition model). **(a)** Path diagram with standardised path coefficients and corresponding standard errors. Only paths with a path coefficient passing a *P*-value threshold of 0.05 are shown. Full information on path coefficients and their standard errors can be found in S3 Table. **(b)** Standardised variance explained by genetic and residual factors modelled in (a). * The proportion of phenotypic variance explained by genetic factor A4 in receptive vocabulary at 38 months is negligible. Abbreviations: CDI, Communicative Development Inventory; Exp, expressive; m, months of age; Rec, receptive; voc, vocabulary

Expressive vocabulary at 24 months loaded on a second genetic factor (A2), explaining an additional 10.1%(SE=4.0%) of the phenotypic variation (path-coefficient a_22_: 0.32(SE=0.06), *P*=4×10^−7^; Fig 2, S3 Table) and the majority of the SNP-h^2^ (factorial co-heritability: 68.8%(SE=23.4%), S4 Table). This genetic factor was also shared with both expressive (path coefficient a_32_:0.27(SE=0.09), *P*=0.005) and receptive (path coefficient a_42_:0.33(SE=0.08), *P*=4×10^−5^) vocabulary at 38 months, accounting for 7.1%(SE=5.0%) and 11.0%(5.3%) of the phenotypic variation, respectively (Fig 2, S3 Table). For receptive vocabulary at 38 months, this genetic factor captured the majority of the SNP-h^2^ (factorial co-heritability: 88.9%(SE=23.1%), Table S4), suggesting a largely shared genetic aetiology with expressive vocabulary at 24 months, as confirmed by their high genetic correlation (GSEM-r_g_=0.78(SE=0.19), Fig 1c).

The third genetic factor (A3) was only related to expressive vocabulary at 38 months (path coefficient a_33_:0.29(SE=0.08), *P*=0.001) and explained 8.2%(SE=4.9%) of the phenotypic variation (Fig 2, S3 Table), corresponding to nearly half of the SNP-h^2^ (factorial co-heritability: 47.0%(SE=25.1%), S4 Table). This genetic factor was unrelated to receptive vocabulary at 38 months (path coefficient a_43_:0.12(SE=0.12), *P*=0.35). Thus, it is likely that the genetic correlation between expressive and receptive vocabulary at 38 months (GSEM-r_g_=0.82(SE=0.12, Fig 1c) is primarily driven by genetic variance shared with expressive vocabulary at 24 months.

Finally, there was little support for the presence of a fourth genetic factor (A4) that would be exclusively related to receptive vocabulary at 38 months (Fig 2, S3 Table). However, according to findings from our previous work, such a factor is likely to account only for very little phenotypic variance in receptive vocabulary at 38 months (15). Therefore, it may only become detectable once modelled together with other heritable traits sharing underlying genetic influences.

### Stage 2: Multivariate genetic variance structures between early-life vocabulary and mid-childhood reading, verbal and performance intelligence

In a second step, we assessed the emergence of genetic links with mid-childhood reading accuracy/comprehension at 7 years, verbal intelligence quotient scores (VIQ) at 8 years and performance intelligence quotient scores (PIQ) at 8 years (S1 Table) across the studied vocabulary measures during the first three years of life, using rank-transformed measures. The selected measures of reading and verbal intelligence are representative of previously reported genetic association patterns between vocabulary at 38 months and a wide spectrum of language, literacy and cognitive abilities in ALSPAC (15). We contrast these verbal abilities with a measure of non-verbal intelligence (PIQ) to evaluate differences in developmental association patterns with respect to early-life vocabulary. Thus, the model from the first step (Fig 2) was extended to include, in turn, each of the three mid-childhood skills, resulting in three further SEMs (with measures included in chronological order).

At the phenotypic level, all early-life vocabulary measures showed low to modest correlations with both mid-childhood verbal and non-verbal skills (Fig 3a), with the largest phenotypic correlation between receptive vocabulary at 38 months and VIQ at 8 years (r_p_=0.26). The selected mid-childhood skills, reading, VIQ and PIQ, were all moderately heritable, with GCTA-SNP-h^2^ estimates of 42%(SE=6%), 54%(SE=7%) and 26%(SE=7%), respectively. These estimates largely corresponded to GSEM-SNP-h^2^ estimates (S2 Table). Using GCTA, bivariate genetic correlations of mid-childhood skills with early-life vocabulary measures (Fig 3b) revealed moderate genetic correlations of VIQ with expressive vocabulary at 24 (GCTA-r_g_=0.41(SE=0.14),*P*=0.003) and 38 months (GCTA-r_g_=0.38(SE=0.14),*P*=0.003), but high genetic correlations of both verbal and non-verbal skills with receptive vocabulary at 38 months (reading GCTA-r_g_=0.83(SE=0.25), *P*=9×10^−6^; VIQ GCTA-r_g_=0.95(SE=0.23), *P*=1×10^−8^; PIQ GCTA-r_g_=0.68(SE=0.28),*P*=0.004). GCTA and GSEM genetic correlation estimates were highly consistent, with overlapping 95%-CIs (Figs 3b and 3c). GSEM-estimated residual correlations between early-life and mid-childhood measures were low (Fig 3d).

**Fig 3.**
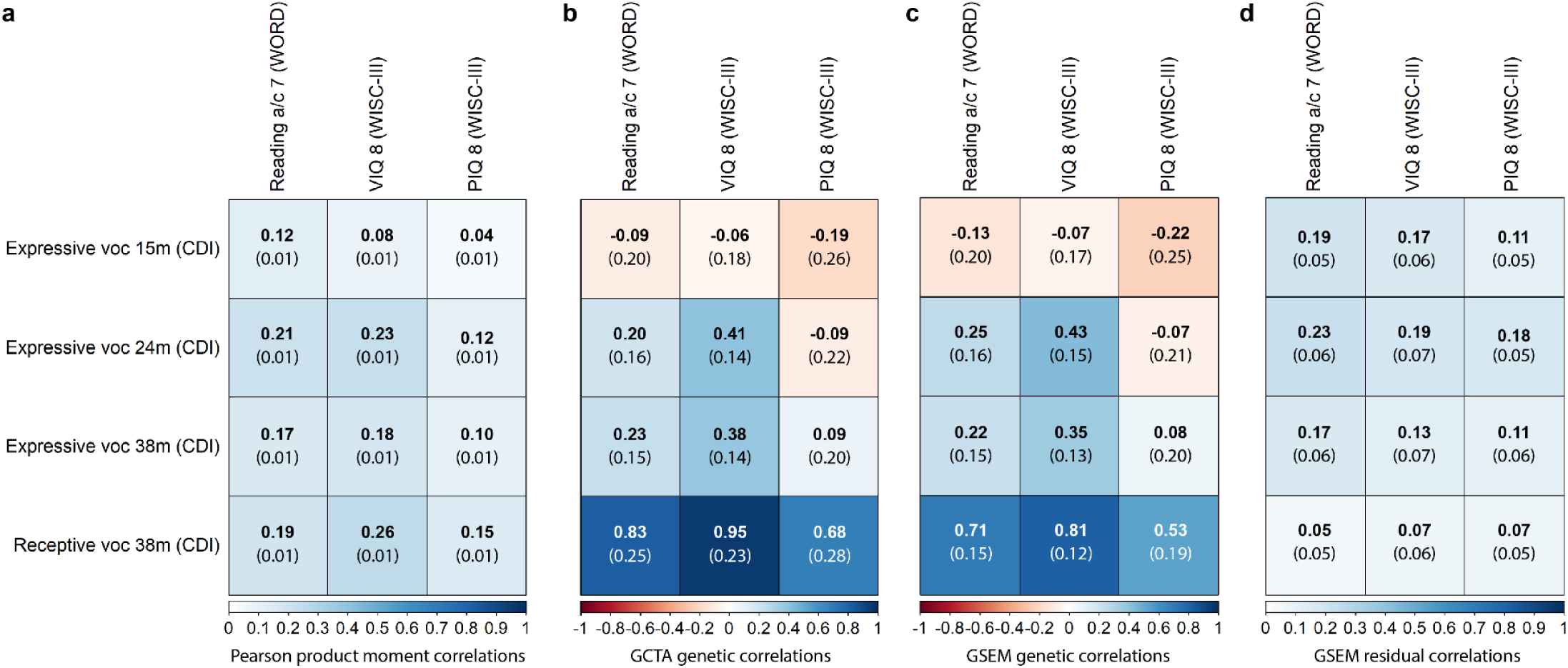
Phenotypic, genetic and residual correlations between early-life vocabulary scores and mid-childhood reading, verbal intelligence and performance intelligence. Correlation patterns are shown for rank-transformed measures with sufficient evidence for SNP-h^2^ (*P*<0.05) Standard errors are shown in brackets. **(a)** Phenotypic correlations were estimated with Pearson correlation coefficients. **(b)** GCTA genetic correlations based on GREML. **(c)** GSEM genetic correlations. **(d)** GSEM residual correlations. Abbreviations: a, accuracy; c, comprehension; CDI, Communicative Development Inventory; GCTA, Genome-based Restricted Maximum Likelihood as implemented in genome-wide complex trait analysis (GCTA) software; GREML, Genome-based restricted maximum likelihood; GSEM, genetic-relationship-matrix structural equation modelling; m, months; PIQ; performance intelligence quotient; VIQ; verbal intelligence quotient, voc, vocabulary; WISC-III, Wechsler Intelligence Scale for Children III; WORD, Wechsler Objective Reading Dimension Genetic analyses were conducted using genetic relationship matrices based on directly genotyped SNPs and individuals with a genetic relationship of <0.05

Using multivariate structural models, our results showed, first, that there is little evidence for genetic links between expressive vocabulary at 15 months (A1) and vocabulary, reading or cognition abilities after the age of 24 months (Fig 4, S6, S7, S8 Tables). Second, the developmentally novel genetic factor emerging for expressive vocabulary at 24 months (A2), explained further genetic variance in receptive and expressive vocabulary at 38 months (as outlined above) and, importantly, mid-childhood verbal skills. Specifically, it was related to both reading accuracy/comprehension (path coefficient a_52_:0.25(SE=0.12), *P*=0.04) and VIQ (path coefficient a_52_=0.42(SE=0.13), *P*=0.001), and accounted for 6.4%(6.2%) and 17.9%(11.1%) of their phenotypic variation, respectively (Fig 4, S6 and S7 Tables). However, this genetic factor was not linked to PIQ at 8 years (path coefficient a_52_:-0.03(SE=0.12), *P*=0.78)(Fig 4e and 4f, S8 Table). These findings may reflect some genetic specificity for verbal skills (reading and VIQ), compared to non-verbal cognition, though the 95%-CIs for the identified path coefficients overlap (path coefficients a_52_-reading accuracy/comprehension: 95%-CI=0.01-0.49, a_52_-VIQ: 95%-CI=0.17-0.68, a_52_-PIQ: 95%-CI=-0.26-0.20, derived assuming normality). Third, genetic influences identified for expressive vocabulary at 38 months (A3) were unrelated to receptive vocabulary assessed at the same age (as outlined above) and later mid-childhood abilities (Fig 4, S6, S7, S8 Tables). Thus, the genetic correlation observed between expressive vocabulary at 38 months and mid-childhood VIQ (GSEM-r_g_=0.35(SE=0.13), Fig 3c) is primarily driven by genetic variance shared with expressive vocabulary at 24 months. Fourth, joint modelling of early-life vocabulary measures with mid-childhood abilities enabled the identification of a genetic factor that affects receptive vocabulary at 38 months (A4) and that is independent of early-life expressive vocabulary genetic factors (path coefficient a_44_:0.15(SE=0.07), *P*=0.04, Fig 4c). Although this genetic factor accounted for only a tiny proportion of the phenotypic variation in receptive vocabulary at 38 months (2.1%(SE=1.9%)), it explained 33.0%(SE=8.2%), 36.1%(SE=11.5%) and 24.7%(SE=7.5%) of the phenotypic variation in reading accuracy/comprehension, VIQ and PIQ, respectively (path coefficients a_54_-reading accuracy/comprehension:0.57(SE=0.07), *P*<1×10^−10^; a_54_-VIQ: 0.60(0.10), *P*=3×10^−10^; a_54_-PIQ: 0.50(0.08), *P*<1×10^−10^). The genetic variance explained by genetic factor A4 corresponds to the majority of the estimated SNP-h^2^ for mid-childhood abilities, as indicated by factorial co-heritabilities (reading: 82.3%(SE=16.1%), VIQ: 66.4%(SE=19.9%), PIQ: 91.8%(SE=15.1%), S9 Table). Finally, there was little evidence for novel genetic factors emerging during mid-childhood (A5, Fig 4), consistent with previous findings (15). Thus, the fitted multivariate models for early-life vocabulary and mid-childhood skills were consistent with both the identified multivariate genetic architecture of early-life vocabulary (Fig 2) and the previously reported amplification of genetic factors for vocabulary at 38 months (15).

**Fig 4.**
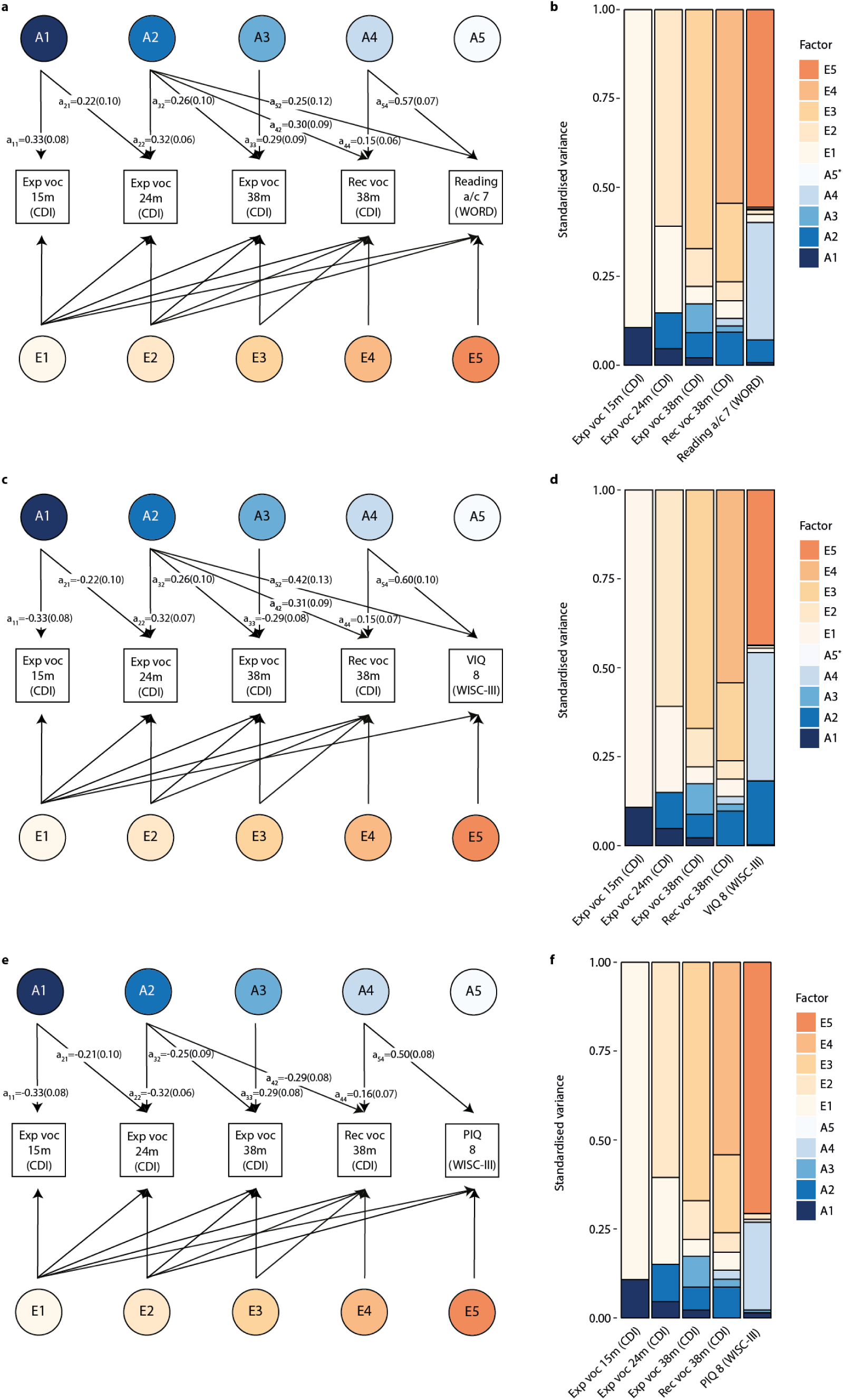
Structural models of early-life vocabulary and mid-childhood reading and cognition. Genetic-relationship matrix structural equation modelling (GSEM) of early-life vocabulary scores (15, 24 and 38 months of age) in combination with mid-childhood **(a,b)** reading accuracy/comprehension at 7 years, **(c,d)** VIQ scores at 8 years or **(e,f)** PIQ scores at 8 years, based on all available observations for children across development (N≤6,524). **(a,c,e)** Path diagrams with standardised path coefficients and corresponding standard errors including mid-childhood **(a)** reading accuracy/comprehension, **(c)** VIQ and **(e)** PIQ outcomes. Only paths with a path coefficient passing a *P*-value threshold of 0.05 are shown. Full information on all path coefficients and their standard errors can be found in S6, S7, S8 Tables. **(b,d,f)** Standardised variance explained by genetic and residual factors as modelled in a,c,e for models including **(b)** reading accuracy/comprehension, **(d)** VIQ, and **(f)** PIQ. * The proportion of phenotypic variance explained by genetic factor A5 is negligible. Abbreviations: a, accuracy; c, comprehension; CDI, Communicative Development Inventory; Exp, expressive; m, months of age; Rec, receptive; PIQ, performance intelligence quotient; VIQ, verbal intelligence quotient; voc, vocabulary; WISC-III, Wechsler Intelligence Scale for Children III; WORD, Wechsler Objective Reading Dimension

The phenotypic covariance of mid-childhood reading, VIQ and PIQ with receptive vocabulary at 38 months (Fig 3a) was primarily due to genetic covariance, with bivariate heritability estimates of 0.87(SE=0.21), 0.88(SE=0.16) and 0.68(SE=0.27), respectively (S10 Table). This is consistent with little evidence for residual correlation between receptive vocabulary at 38 months and mid-childhood measures (Fig 3d). For verbal mid-childhood skills, such as VIQ, evidence for bivariate heritability with early-life expressive vocabulary was already detectable at 24 months of age (bivariate heritability: 0.54(SE=0.19)), as well as at 38 months of age (bivariate heritability: 0.60(SE=0.24)).

## Discussion

This genome-wide longitudinal analysis of vocabulary size during the first three years of life assessed in unrelated children demonstrates that the genetic architecture underlying expressive and receptive vocabulary is dynamic, with evidence for both age- and ability-specific genetic influences. Genetic continuity was found for two independent early-life genetic factors, which contribute to the genetic variance of reading and cognitive skills in mid-childhood. One stable early-life genetic source of variation was related to expressive vocabulary and emerged at 24 months of age, accounting for between 6.4% and 17.9% of the phenotypic variation in mid-childhood abilities, especially verbal skills such as reading and VIQ. A second, independent and stable early-life genetic factor was identified for receptive vocabulary at 38 months and explained between 24.7% and 36.1% of the phenotypic variance in both mid-childhood verbal and non-verbal cognitive abilities, including PIQ, corresponding to the majority of SNP-h^2^ (≥66%). Given the modest SNP-h^2^ of early-life vocabulary scores, ranging from 11% to 18%, this suggests not only genetic stability, but also an amplification of early genetic variance during the life-course that contributes to the markedly increased SNP-h^2^ of later-life reading and cognition (27% to 54%).

The identification of multiple independent genetic factors related to vocabulary during the first three years of life may reflect rapid changes in mastering behavioral and language skills. Genetic influences identified for expressive vocabulary at 15 months (A1) were also related to expressive vocabulary at 24 months, but were not linked to vocabulary, reading or cognition measures beyond this age. Thus, these early genetic influences might primarily affect the very first stages of language development that, once achieved, have little impact on subsequent verbal and cognitive development. A plausible candidate process for this is the acquisition of phonological skills to identify phonemes and sequences from speech and their storage for future production (23).

The stable independent genetic factor emerging for expressive vocabulary at 24 months (A2) contributes to the genetic architectures underlying verbal processes throughout childhood, in contrast to genetic factor A1. Specifically, the genetic influences captured by A2 were related to both expressive and receptive vocabulary at 38 months, as well as mid-childhood verbal abilities such as reading and VIQ, but not PIQ (although 95%-CIs of estimated genetic path coefficients overlap with those for PIQ). This genetic factor may reflect stages of language learning that take place after the production of words in isolation at the age of 10 to 15 months (3). This includes, for example, an increasing vocabulary size as well as the use of more complex grammatical structures, marked by the emergence of two-word combinations around the age of 18 to 24 months (1,24). It has been shown that lexical and grammatical development share underlying acquisition mechanisms (25) and measures of expressive vocabulary and grammatical development at two and three years of age are both phenotypically and genetically correlated (11).

Expressive vocabulary at 38 months loaded on an additional independent genetic factor (A3) that was not related to receptive vocabulary at the same age, nor to any of the studied mid-childhood reading and IQ measures. This genetic factor may, thus, involve genetic associations with processes that affect expressive vocabulary at an early age, but do not play a role in later cognition. They may, for example, reflect social abilities, which are known to impact on vocabulary development and vice versa (26). Note that expressive vocabulary at 38 months is nonetheless genetically related to mid-childhood verbal processes due to shared genetic influences that were already detectable at 24 months (A2).

The majority of SNP-h^2^ for mid-childhood reading, VIQ and also PIQ was accounted for by a genetic factor that emerged at 38 months of age for receptive vocabulary (A4), consistent with previous findings (15). Although this stable genetic factor explained only a very small part of the phenotypic variance in receptive vocabulary (2.1%), it accounted from 66% to 92% of the phenotypic variation in later reading performance, verbal and non-verbal cognition, with very little residual contributions. Due to the wide spectrum of associated mid-childhood phenotypes that are linked with this genetic factor, including both later verbal and non-verbal cognitive abilities, it is possible that the genetically encoded biological processes are important for cognitive development in general. It merits noting that the genetic factor A4 was only detectable once modelled together with a mid-childhood skill sharing underlying genetic variance, probably due to the low proportion of phenotypic variation that it explained in early-life receptive vocabulary.

Previous twin studies demonstrating genetic links between language use in early childhood and later language/literacy skills have been based on a latent factor approach jointly capturing genetic variance of expressive language skills between the ages of 2 and 4 years (16,17). Here, we used a sample of unrelated children with genome-wide genotyping data and distinguish language measures during the first three years of life based on both modality and age at assessment. We extend and refine the previous twin findings by showing that (i) early-life expressive vocabulary at 15 months of age is influenced by a genetic factor that is only shared across expressive vocabulary scores during infancy, and (ii) that there are at least two independent genetic factors during early life that are associated with mid-childhood reading and cognition. Genetic associations with mid-childhood verbal cognitive processes arise as early as 24 months of age, whereas genetic influences that are relevant for mid-childhood general cognitive development emerge as early as 38 months of age for receptive vocabulary, and are independent of expressive vocabulary. This latter distinction is important as receptive vocabulary at 38 months also shares a genetic factor with expressive vocabulary at 24 months and subsequent reading and verbal intelligence. The diversity in genetic factors may implicate differences in overarching cognitive processes that are already detectable during toddlerhood. This is important as genetic influences associated with early-life vocabulary could fully account for the SNP-h^2^ of mid-childhood reading, verbal and non-verbal intelligence. The presence of such genetic stability implicating verbal processes and general cognition from toddlerhood to, at least, mid-childhood may, furthermore, suggests shared biological underpinnings. Thus, joint genome-wide association study analyses across developmental stages may facilitate an increase in study power.

In addition to the strengths of this study described above, this study benefits from modelling multivariate genetic variance structures in unrelated individuals directly, based on genome-wide information, using novel structural equation modelling techniques. It is, however, not possible to infer biological mechanisms underlying the identified genetic factor structures with the current methodology. We furthermore exploit the phenotypic richness of the ALSPAC cohort, including longitudinally assessed vocabulary measures during early development as well as reading and cognitive outcomes in mid-childhood. This study has also several limitations. Given the rapidly changing nature of early vocabulary size, increasingly larger and complex word lists are required to reliably assess vocabulary size at 24 and 38 months compared to 15 months of age. Thus, the observed differences in genetic factor structures during early life may reflect differences in CDI instruments, although this is unlikely to fully explain our findings, given substantial phenotypic correlations between expressive vocabulary scores at 15 and 24 months of age (r_p_=0.53). Furthermore, vocabulary assessments at 38 months of age might be affected by ceiling effects, as the MacArthur CDI:Words & Sentences was developed for children up to 30 months (5). This may have reduced phenotypic variation and, thus, power to detect genetic variance components at 38 months. In addition, it has recently been shown that heritability and genetic relationships estimated in samples of unrelated individuals, especially for cognition-related traits (27,28), might be inflated by indirect genetic effects, reflecting a type of gene-environment correlation (29). The observed association patterns between early-life vocabulary and mid-childhood reading and cognitive skills may therefore represent both shared genetic variance and indirect genetic effects. Future research using family-based data is warranted to assess the impact of indirect genetic effects on the reported association patterns. Finally, the sparcity of large data sets with longitudinal information on expressive and receptive vocabulary during infancy and toddlerhood, in addition to genome-wide data, currently prevents a direct replication of our findings in independent cohorts.

Taken together, our findings reveal a dynamic genetic landscape underlying vocabulary during the first three years of life. We found evidence for genetic continuity of two independent early-life genetic factors that contribute to both verbal and general cognitive abilities in mid-childhood and manifest at different developmental stages during early-life language development. Thus, the genetic foundations for both mid-childhood reading and cognition lie in toddlerhood, but are diverse, and may implicate aetiological differences in overarching cognitive processes that are detectable long before the age of schooling.

## Methods

### Sample description and trait selection

#### Cohort information

Participants were born in 1991 or 1992 and included in ALSPAC, a UK population-based birth cohort (S1 Appendix)(18,19). Ethical approval was provided by the ALSPAC Ethics and Law Committee and the Local Research Ethics Committees. Informed consent for questionnaire and clinical data was obtained from participants following recommendations of the ALSPAC Ethics and Law Committee at the time. Consent for biological samples was collected in accordance with the Human Tissue Act (2004).

#### Genetic analyses

Genotyping and genotype calling was performed using the Illumina HumanHap550 quad chip and Illumina GenomeStudio software. Quality control of genetic data was applied using PLINK (v1.07)(30) at both the SNP and individual level following standard procedures. Individuals were excluded in case of gender mismatch between reported and genetic sex information, >3% missing SNP information, non-European ancestry, or interindividual relatedness (genomic relatedness>0.05). SNPs were excluded if they had a low call rate (<99%), were rare (<1%) and/or deviated from Hardy-Weinberg equilibrium (*P*<5×10^−7^). After quality control, 7,924 children and 465,740 SNPs with high-quality genetic data remained.

#### Early-life vocabulary measures

Expressive and receptive vocabulary was assessed at 15, 24 and 38 months of age using parental-reports (predominantly mother) of age-specific defined word lists adapted from the MacArthur Communicative Development Inventory (CDI). At 15 months, expressive and receptive vocabulary were assessed with an abbreviated version of the MacArthur CDI:Words & Gestures (133 words, 8 to 16 months of age)(31). Scores were recorded as the number of words a child could “say and understand” (expressive vocabulary), and “understand” plus “say and understand” (receptive vocabulary), respectively. At 24 and 38 months of age, an abbreviated vocabulary list from the MacArthur CDI:Words & Sentences (123 words, 16-30 months of age)(5) was used. At both ages, expressive vocabulary was ascertained as the total number of words a child could “say” plus “say and understand”. Receptive vocabulary at 38 months was measured as the total number of words a child could “understand” plus “say and understand”. The receptive vocabulary score at 24 months was excluded due to discrepancies in the applied coding scheme (reflecting the total number of words a child could “understand” only, excluding words a child could “say and understand”), and recoded scores have not yet been released by ALSPAC.

CDI expressive vocabulary scores have high reliability and validity, showing correlations with direct assessments of over 0.70 (32,33). Receptive vocabulary assessed using parental report correlated 0.55 with direct assessment (33). In total, N≤6,524 children (Table 1) had vocabulary scores and genome-wide genetic data available for analyses.

#### Mid-childhood measures

For the selection of mid-childhood measures, we build on our previous work identifying genetic links between vocabulary at 38 months and thirteen mid-childhood/adolescent literacy and cognitive measures (15). As it is not possible, due to computational constrains, to study longitudinal genetic architectures of early-life vocabulary measures in combination with a wide spectrum of mid-childhood language, literacy and cognitive abilities, we selected three mid-childhood measures that are representative of previously observed developmental association patterns (15) (N≤5,296; S1 Table). The studied mid-childhood measures included reading accuracy/comprehension at 7 years, assessed using the Wechsler Objective Reading Dimensions (WORD)(34), as well as both VIQ and PIQ assessed at 8 years using the Wechsler Intelligence Scale for Children (WISC-III)(35). Detailed descriptions, including validity and reliability, of each measure are available in the Supporting Information (S2 Appendix).

#### Phenotype transformation

All early-life vocabulary and mid-childhood measures were adjusted for sex, age (except for VIQ and PIQ as they were derived using age-specific norms), and the first two principal components (adjusting for subtle differences in ancestry (36)), and subsequently rank-transformed. In addition, early-life vocabulary measures were adjusted for age squared, as vocabulary develops rapidly during early childhood (37). Phenotypic correlations between early-life vocabulary measures were estimated using untransformed (Spearman rank-correlation) and rank-transformed (Pearson correlation) scores respectively, and patterns were largely unaffected by trait transformation (S1 Fig). Phenotypic correlations between early-life vocabulary and mid-childhood reading, VIQ and PIQ measures were estimated using rank-transformed (Pearson correlation) scores only.

### Genome-wide Complex Trait Analysis

Total SNP-h^2^ was estimated using Genome-based restricted maximum likelihood (GREML) analyses (38,39), as implemented in GCTA software (21), based on a GRM including directly genotyped SNPs only (GCTA-SNP-h^2^). Measures with little evidence for GCTA-SNP-h^2^ (*P*>0.05) were excluded from further analyses.

Bivariate GREML (39) was applied to estimate bivariate genetic correlations among early-life vocabulary measures and between early-life vocabulary and mid-childhood reading, VIQ and PIQ measures.

### Multivariate genetic analyses

To study the genetic architecture of vocabulary in a developmental context, we used Genetic-relationship-matrix Structural Equation Models (GSEMs)(20). This is a multivariate structural equation modelling technique, which combines multivariate analysis methodologies established in twin research (40,41) with estimates of genetic relationships between unrelated individuals, as captured by genome-wide genetic markers (20) (S3 Appendix). Specifically, GSEMs dissect the phenotypic covariance structure into one or more additive genetic factors (A), capturing genetic variance tagged by common genotyped SNPs, as well as one or more residual factors (E) that resemble the residual variance, containing both untagged genetic variation and unique environmental influences (including measurement error). Here, multivariate GSEMs were fitted to the data through a Cholesky decomposition model, with the phenotypic variance decomposed into as many latent genetic and residuals factors as there are observed variables, without any restrictions on the structure (42) (S3 Appendix). Structural models were based on all available observations across individuals and thus allow for missing data (saturated model; R:gsem library, version 0.1.5). Genetic relationships between individuals were assessed with GRMs, including directly genotyped SNPs only, as implemented in GCTA software (21).

## Acknowledgements

We are extremely grateful to all the families who took part in this study, the midwives for their help in recruiting them, and the whole ALSPAC team, which includes interviewers, computer and laboratory technicians, clerical workers, research scientists, volunteers, managers, receptionists and nurses. This publication is the work of the authors and EV and BSTP will serve as guarantors for the contents of this paper.

## Data availability statement

Information about ALSPAC data is available through a fully searchable data dictionary (http://www.bris.ac.uk/alspac/researchers/data-access/data-dictionary/). Access to ALSPAC data can be obtained as described within the ALSPAC data access policy (http://www.bristol.ac.uk/alspac/researchers/access/). All analyses were performed using freely accessible software. Requests for scripts or other analysis details can be sent via email to the corresponding author.

## Authors’ contribution statement

BSTP developed the study concept and EV contributed to the study design. EV performed the data analysis and interpretation under the supervision of BSTP. EV and BSTP drafted the manuscript and CYS, SEF and PSD provided critical revisions. All authors approved the final version of the manuscript for submission.

## Supporting information

S1 Appendix. ALSPAC description

S2 Appendix. Mid-childhood ALSPAC measures

S3 Appendix. Genetic-relatedness-matrix Structural equation modelling

S4 Appendix. Factorial co-heritability

S5 Appendix. Bivariate heritability

S6 Appendix. Websites

S1 Table. Mid-childhood measures in the Avon Longitudinal Study of Parents and Children

S2 Table. SNP heritability estimates

S3 Table. Standardised path coefficients and variance explained for early-life vocabulary measures

S4 Table. Factorial co-heritability for early-life vocabulary measures

S5 Table. Bivariate heritability for early-life vocabulary measures

S6 Table. Standardised path coefficients and variance explained for early-life vocabulary and mid-childhood reading accuracy/comprehension

S7 Table. Standardised path coefficients and variance explained for early-life vocabulary and mid-childhood verbal intelligence

S8 Table. Standardised path coefficients and variance explained for early-life vocabulary and mid-childhood performance intelligence

S9 Table. Factorial co-heritability for genetic factors contributing to mid-childhood reading, verbal intelligence and performance intelligence

S10 Table. Bivariate heritability for early-life vocabulary measures and mid-childhood reading, verbal intelligence and performance intelligence

S1 Fig: Phenotypic correlations among early-life vocabulary measures

S2 Fig: Path diagram for a trivariate trait

## References

1. Fenson L, Dale PS, Reznick JS, Bates E, Thal DJ, Pethick SJ. Variability in early communicative development. Monogr Soc Res Child Dev. 1994;59(5):1–173; discussion 174–85.

2. Kennison SM. Introduction to language development. 1st ed. SAGA; 2014. 469 p.

3. Clark EV. First language acquisition. New York: Cambridge University Press; 2016.

4. Goldfield BA, Reznick JS. Early lexical acquisition: rate, content, and the vocabulary spurt. J Child Lang. 1990 Feb;17(1):171–83.

5. Fenson L, Dale P, Reznick JS, Thal D, Bates E, Hartung J, et al. User’s Guide and Technical Manual for the MacArthur Communicative Development Inventories. San Diego: Singular Publishing; 1993.

6. Burger A, Chong I. Receptive Vocabulary. In: Goldstein S, Naglieri JA, editors. Encyclopedia of Child Behavior and Development. Boston, MA: Springer US; 2011. p. 1231–1231.

7. Owens Jr. R E. Language Development: An Introduction. 9 edition. Pearson; 2015. 462 p.

8. Bergelson E, Swingley D. At 6-9 months, human infants know the meanings of many common nouns. Proc Natl Acad Sci USA. 2012 Feb 28;109(9):3253–8.

9. Fenson L, Marchman VA. MacArthur-Bates Communicative Development Inventories: User’s Guide and Technical Manual. Paul H. Brookes Publishing Company; 2007. 188 p.

10. Frank MC, Braginsky M, Yurovsky D, Marchman VA. Wordbank: an open repository for developmental vocabulary data. J Child Lang. 2017 May;44(3):677–94.

11. Dionne G, Dale PS, Boivin M, Plomin R. Genetic Evidence for Bidirectional Effects of Early Lexical and Grammatical Development. Child Development. 2003;74(2):394–412.

12. Dale PS, Dionne G, Eley TC, Plomin R. Lexical and grammatical development: a behavioural genetic perspective. Journal of Child Language. 2000 Oct;27(03):619–642.

13. Reznick JS, Corley R, Robinson J. A longitudinal twin study of intelligence in the second year. Monogr Soc Res Child Dev. 1997;62(1):i–vi, 1–154; discussion 155-60.

14. St Pourcain B, Cents RA, Whitehouse AJ, Haworth CM, Davis OS, O’Reilly PF, et al. Common variation near ROBO2 is associated with expressive vocabulary in infancy. Nature communications. 2014;5:4831.

15. Verhoef E, Shapland CY, Fisher SE, Dale PS, Pourcain BS. The developmental origins of genetic factors influencing language and literacy: Associations with early-childhood vocabulary. Journal of Child Psychology and Psychiatry [Internet]. 2020 Sep 14; Available from: https://doi.org/10.1111/jcpp.13327

16. Harlaar N, Hayiou-Thomas ME, Dale PS, Plomin R. Why Do Preschool Language Abilities Correlate With Later Reading? A Twin Study. J Speech Lang Hear Res. 2008 Jun 1;51(3):688–705.

17. Hayiou-Thomas ME, Dale PS, Plomin R. The etiology of variation in language skills changes with development: a longitudinal twin study of language from 2 to 12 years. Dev Sci. 2012 Mar;15(2):233–49.

18. Fraser A, Macdonald-Wallis C, Tilling K, Boyd A, Golding J, Davey Smith G, et al. Cohort Profile: the Avon Longitudinal Study of Parents and Children: ALSPAC mothers cohort. Int J Epidemiol. 2013 Feb;42(1):97–110.

19. Boyd A, Golding J, Macleod J, Lawlor DA, Fraser A, Henderson J, et al. Cohort Profile: the ‘children of the 90s’--the index offspring of the Avon Longitudinal Study of Parents and Children. Int J Epidemiol. 2013 Feb;42(1):111–27.

20. St Pourcain B, Eaves LJ, Ring SM, Fisher SE, Medland S, Evans DM, et al. Developmental changes within the genetic architecture of social communication behaviour: A multivariate study of genetic variance in unrelated individuals. Biological Psychiatry. 2017 Sep 28;83:598–606.

21. Yang J, Lee SH, Goddard ME, Visscher PM. GCTA: a tool for genome-wide complex trait analysis. Am J Hum Genet. 2011 Jan 7;88(1):76–82.

22. Fenson L, Dale PS, Reznic S. Technical Manual for the MacArthur Communicative Development Inventories. Developmental Psychology Laboratory; 1991.

23. Curtin S, Archer SL. Speech perception. In: Bavin EL, Naigles LR, editors. The Cambridge Handbook of Child Language. 2nd ed. Cambridge: Cambridge University Press; 2015. p. 137–58. (Cambridge Handbooks in Language and Linguistics).

24. Hoff E. Language Development. Cengage Learning; 2013. 482 p.

25. Bates E, Goodman JC. On the inseparability of grammar and the lexicon: Evidence from acquisition. In: Language development: The essential readings. Malden: Blackwell Publishing; 2001. p. 134–62. (Essential readings in developmental psychology).

26. Silverman RD, Hartranft AM. Developing Vocabulary and Oral Language in Young Children. Guilford Publications; 2014. 274 p.

27. Selzam S, Ritchie SJ, Pingault J-B, Reynolds CA, O’Reilly PF, Plomin R. Comparing Within- and Between-Family Polygenic Score Prediction. The American Journal of Human Genetics. 2019 Aug 1;105(2):351–63.

28. Cheesman R, Hunjan A, Coleman JRI, Ahmadzadeh Y, Plomin R, McAdams TA, et al. Comparison of Adopted and Nonadopted Individuals Reveals Gene–Environment Interplay for Education in the UK Biobank: Psychological Science [Internet]. 2020 Apr 17; Available from: https://journals.sagepub.com/doi/full/10.1177/0956797620904450

29. Kong A, Thorleifsson G, Frigge ML, Vilhjalmsson BJ, Young AI, Thorgeirsson TE, et al. The nature of nurture: Effects of parental genotypes. Science. 2018 Jan 26;359(6374):424–8.

30. Purcell S, Neale B, Todd-Brown K, Thomas L, Ferreira MAR, Bender D, et al. PLINK: A Tool Set for Whole-Genome Association and Population-Based Linkage Analyses. Am J Hum Genet. 2007 Sep;81(3):559–75.

31. Fenson L, Pethick S, Renda C, Cox JL, Dale PS, Reznick JS. Short-form versions of the MacArthur Communicative Development Inventories. Applied Psycholinguistics. 2000 Mar;21(1):95–116.

32. Dale PS. The validity of a parent report measure of vocabulary and syntax at 24 months. J Speech Hear Res. 1991 Jun;34(3):565–71.

33. Ring ED, Fenson L. The correspondence between parent report and child performance for receptive and expressive vocabulary beyond infancy. First Language. 2000 Jun 1;20(59):141–59.

34. WORD, Wechsler Objective Reading Dimensions Manual. Psychological Corporation; 1993. 146 p.

35. Wechsler D, Golombok S, Rust J. WISC-III UK Wechsler Intelligence Scale for Children – Third Edition UK Manual. Sidcup, UK: The Psychological Corporation; 1992.

36. Price AL, Patterson NJ, Plenge RM, Weinblatt ME, Shadick NA, Reich D. Principal components analysis corrects for stratification in genome-wide association studies. Nature Genetics. 2006 Aug;38(8):904–9.

37. Brooks R, Meltzoff AN. Infant gaze following and pointing predict accelerated vocabulary growth through two years of age: a longitudinal, growth curve modeling study. J Child Lang. 2008 Feb;35(1):207–20.

38. Yang J, Benyamin B, McEvoy BP, Gordon S, Henders AK, Nyholt DR, et al. Common SNPs explain a large proportion of the heritability for human height. Nat Genet. 2010 Jul;42(7):565–9.

39. Lee SH, Yang J, Goddard ME, Visscher PM, Wray NR. Estimation of pleiotropy between complex diseases using single-nucleotide polymorphism-derived genomic relationships and restricted maximum likelihood. Bioinformatics. 2012 Oct 1;28(19):2540–2.

40. Martin NG, Eaves LJ. The genetical analysis of covariance structure. Heredity (Edinb). 1977 Feb;38(1):79–95.

41. Neale M, Maes HHM. Methodology for genetic studies of twins and families. Dordrecht: Kluwer Academic Publisher; 2004.

42. Neale M, Boker S, Xie G, Maes HHM. Mx: Statistical modeling. 7th ed. Richmond: Department of Psychiatry; 2006.

